# Gut microbiota influences foraging onset without affecting division of labor and associated physiological hallmarks in honeybees

**DOI:** 10.1101/2023.12.20.570781

**Authors:** Joanito Liberti, Erik T. Frank, Tomas Kay, Lucie Kesner, Maverick Monié--Ibanes, Andrew Quinn, Thomas Schmitt, Laurent Keller, Philipp Engel

**Affiliations:** Department of Ecology and Evolution, University of Lausanne, Switzerland; Department of Fundamental Microbiology, University of Lausanne, Switzerland; Department of Animal Ecology and Tropical Biology, University of Würzburg, Germany

**Keywords:** *Apis mellifera*, behavioral maturation, social behavior, behavioral development, microorganisms, symbiosis

## Abstract

Gut microbes can impact cognition and behavior, but whether they regulate division of labor in animal societies is unknown. We addressed this question using honeybees since they exhibit division of labor between nurses and foragers and because their gut microbiota can be manipulated. Using automated behavioral tracking and controlling for co-housing effects, we show that gut microbes influence the age at which bees start foraging but have no effects on the time spent in a foraging area and number of foraging trips. Moreover, the gut microbiota did not influence hallmarks of behavioral maturation such as body weight, cuticular hydrocarbon (CHC) profile, hypopharyngeal gland size, and the proportion of bees maturing into foragers. Overall, this study shows that the honeybee gut microbiota does not affect division of labor but rather plays an important function in controlling the onset of bee foraging.

## Introduction

The relationship between the gut microbiota and the physiology and consequent behavior of animal hosts is of fundamental importance to evolutionary biology and of great applied relevance to animal agriculture and human welfare. Beyond the regulation of nutritional intake and immunity, the gut microbiota is a significant determinant of cognition, affecting sensory and social behavior (1–8). Gut microbes can metabolically influence host behavior directly, by producing neuroactive compounds, and indirectly, by releasing secondary products of digestion that interact with the nervous or endocrine system (1, 9). Most studies documenting a link between the microbiota and behavior have focused on the expression of prototypical behaviors at specific stages in an animal’s life. However, behavior can change dramatically with (st)age, and some species even exhibit transitions and reversals between distinct behavioral states. In rodent models it was shown that gut bacteria can influence both the early canalization of behavioral development, with widespread consequences on cognitive ability later in life (10, 11), and the physiological mechanisms that determine behavioral variation within social groups, such as dominance hierarchies (12). So far, few studies have attempted to map the influence of symbiotic organisms onto developmental axes of behavior.

Eusocial insects (ants, termites, some bees and wasps) live in complex societies in which individuals specialize on different tasks during adult life. Morphologically distinct queen and worker ‘castes’ are typically determined early during development and their developmental programs cannot be reversed (13). However, adult workers can sometimes transition between defined physiological / behavioral states (i.e., polyethism) (14, 15). Eusocial insects are therefore studied to understand how morphological, physiological, and behavioral diversity can derive from the same genetic makeup (16). While developmental trajectories are known to be regulated by (epi)genetic mechanisms in response to dietary and environmental cues (17, 18), individuals from different (sub-)castes often show differences in gut microbiota composition or structure (19–23). These differences are generally assumed to be a consequence of different host physiology or dietary preferences. However, whether the gut microbiota could in turn play a regulatory role in division of labor remains unknown (24, 25).

Among eusocial insects, the honeybee has emerged as a model to address these questions (24, 26) because (i) it has a well-characterized, simple and stable ‘core’ gut microbiota (27), (ii) individuals are sterile upon adult emergence, allowing the manipulation of microbiota composition without antibiotic treatment (26), and (iii) it is highly social, exhibiting behaviors that the gut microbiota may influence. The gut microbiota of worker honeybees has been suggested to influence various host phenotypes, including aspects of neurophysiology and consequent cognitive abilities (7, 28–30), collective behavior (7), weight gain (31), and cuticular hydrocarbon (CHC) profiles (32), which are used in nestmate recognition and to indicate behavioral sub-caste (33). However, these phenotypes all covary with behavioral state. Honeybee workers generally spend their first two to three weeks caring for brood inside the hive (‘nursing’) and performing other in-hive tasks. They then undergo a rapid behavioral transition to foraging - regularly leaving the nest in search of food. This transition is regulated by hormones and is associated with profound physiological and behavioral changes, including in CHC profile, weight, gene expression, dietary preference, and gut microbiota composition (21, 23, 34–37). Consequently, it is possible that the detected effects of the gut microbiota on different aspects of honeybee physiology are indirect and mediated by an effect of the gut microbiota on behavioral maturation. For example, all documented effects would be expected if the gut microbiota accelerated or retarded behavioral maturation.

Here we conducted a series of experiments to assess the effect of the gut microbiota on behavioral maturation. We address this at the behavioral level with an automated tracking system in the laboratory, calculating the age at which bees made the first trip to a foraging arena, the proportion of time they spent in the arena, and the total number of foraging trips performed. We also measured several physiological hallmarks of behavioral maturation, such as CHC profile, weight, hypopharyngeal gland size (these glands degenerate during maturation (38)), and gene expression. Overall, our results suggest an effect of the gut microbiota on the timing of the first foraging trip, but not on any of the other maturation-related behaviors or associated physiological hallmarks. This is in contrast to previous studies which suggested that the honeybee gut microbiota modifies the host CHC profile with consequences on nestmate recognition (32), and promotes host weight gain (31). A possible explanation for these discrepancies may be that previous studies used several individuals from the same cage for statistical analyses. Individuals within a cage engage in social interactions and hence they are not independent from each other in aspects of behavior and physiology. Treating them as individual data points in statistical analyses can result in spurious associations between gut microbiota composition and host phenotypes (39, 40).

## Results

### The gut microbiota accelerates the onset of foraging-like behavior under an automated behavioral tracking system

To determine whether the gut microbiota influences the rate of foraging, we reanalyzed behavioral tracking data from a previous study (7). This experiment comprised nine pairs of microbiota-depleted (MD) and microbiota-colonized (CL) sub-colonies consisting of ca. 100 age-matched workers. These bees had been manually extracted from nine hives at the pupal stage and incubated under sterile conditions. The newly emerged adult bees were then inoculated (CL), or not (MD), with a gut homogenate from five nurse bees. Each sub-colony could freely move between a nest-box (30 **°**C, 70% RH in constant darkness) and a foraging arena subject to cycles of light, temperature and humidity mirroring the external environment. The position and orientation of each bee in each sub-colony was tracked by a pair of camera systems using unique matrix barcodes (ARTag library; (41)) attached to the bees’ thoraces. Bees were tracked for a week, starting three days after adult emergence and treatment inoculation, so that the gut microbiota would have fully established (27, 42). There was no significant effect of the microbiota status on the number of trips to the foraging arena (Fig. 1A; Wilcoxon matched-pairs signed-rank test: *V*=29, *P*=0.50) nor the proportion of time spent in the foraging arena (Fig. 1B; Wilcoxon matched-pairs signed-rank test: *V*=25, *P*=0.82). However, CL bees started to perform trips to the foraging arena on average 15 h earlier than MD bees (when bees were between 5-6 days old; Fig. 1C; paired *t*-test: *t*=-4.21, df=8, *P*=0.003). This acceleration of the average age at first foraging occurred in all nine sub-colony pairs.

**Figure 1.**
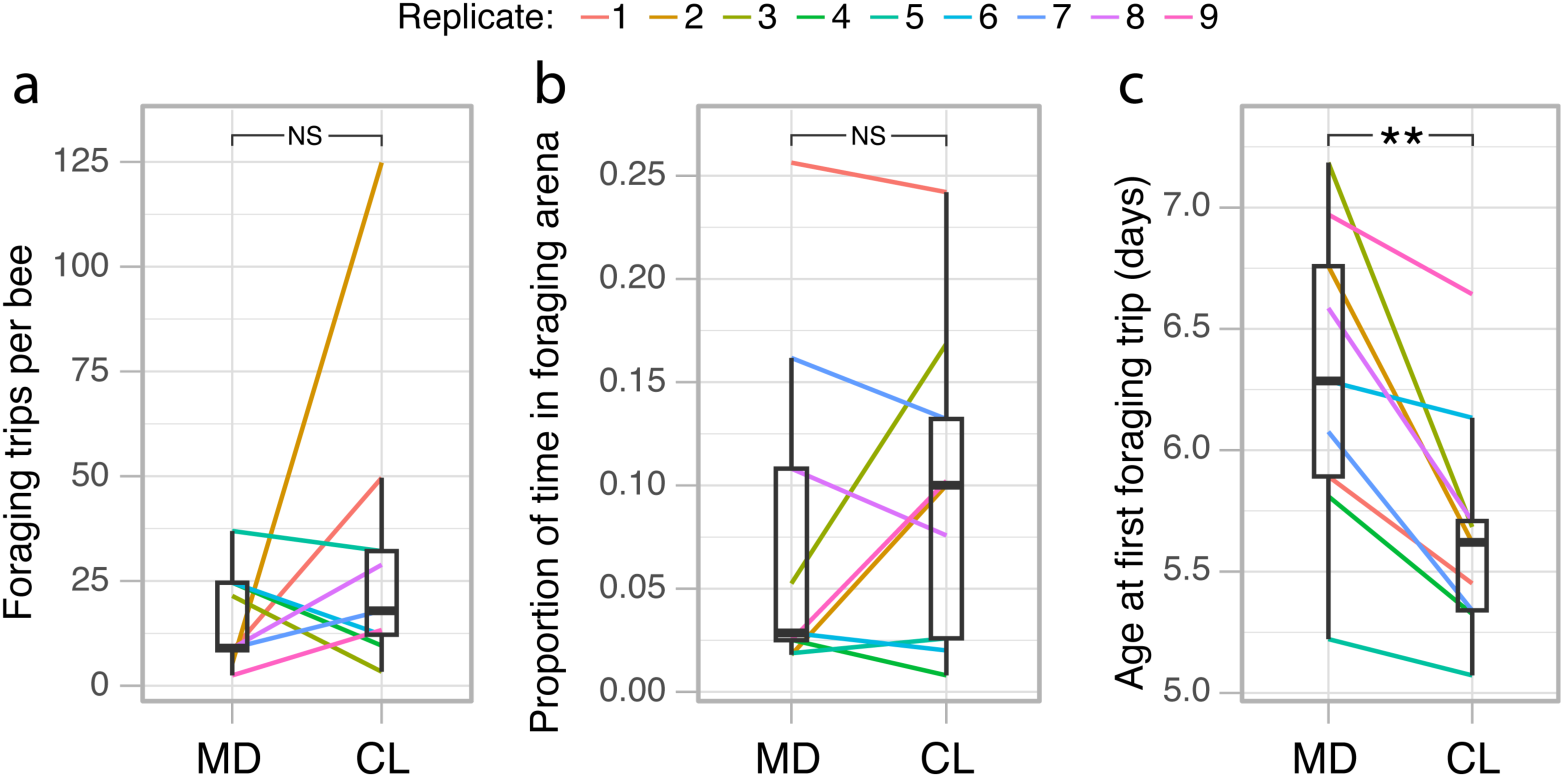
The gut microbiota accelerates the onset of foraging behavior under an automated behavioral tracking system. (a) Average number of trips between the nest and the foraging arena per bee for each sub-colony in the automated behavioral tracking experiment. (b) Average proportion of time spent in the foraging arena per bee for each sub-colony. (c) Average age at which bees made their first trip to the foraging arena for each sub-colony. Lines connect paired sub-colonies and are colored by experimental replicate. Boxplots show the median and first and third quartiles. Whiskers show the extremal values within 1.5× the interquartile ranges above the 75th and below the 25th percentile. \*\**P* < 0.01; NS, not significant, as calculated by paired *t*-test (two-sided).

### The gut microbiota does not modify the CHC profiles of honeybees

The CHC profile of bees changes during the transition from nursing to foraging (37). To assess the effect of the gut microbiota on the CHC profile of bees, we randomly sampled 8-10 bees from each of the 18 sub-colonies (n=177) at the end of the automated behavioral tracking experiment (when bees were 10-day-old) for CHC analysis. Amplicon-sequencing and qPCR analyses targeting the 16S rRNA gene from gut samples of these same bees confirmed that CL and MD bees differed, as expected, in both gut microbiota composition and total load (see Extended Figure 1 in ref. (7)). However, in contrast to the previous study (32), there was no significant effect of the gut microbiota on the CHC profile (Fig. 2A and Supplementary Table 1; permutational multivariate analysis of variance (PERMANOVA) using Bray-Curtis dissimilarities calculated from the centroids of each sub-colony: n=18, *F*(*_1,17_*)=0.89, R^2^=0.04, *P*=0.63).

**Figure 2.**
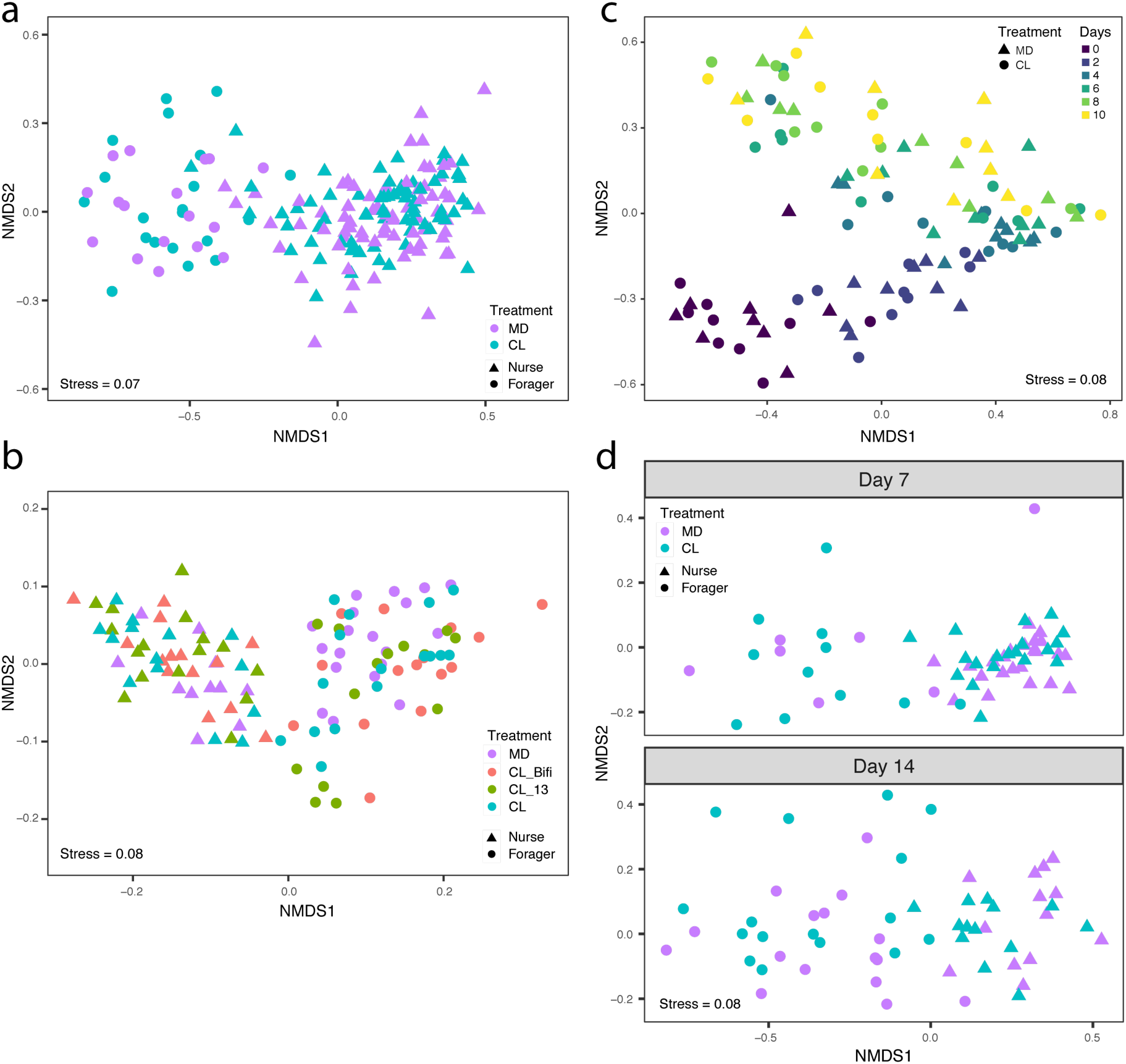
The gut microbiota does not affect CHC profile. (a) Non-metric multidimensional scaling (NMDS) of Bray-Curtis dissimilarities between CHC profiles in the automated behavioral tracking experiment (MD, n=88; CL, n=89). (b) NMDS of Euclidean distances between CHC profiles in the RNA-sequencing experiment, after removal of batch effects from two separate GC-MS runs (MD, n=31; CL_Bifi, n=29; CL_13, n=30; CL, n=30). (c) NMDS of Bray-Curtis dissimilarities between CHC profiles in the longitudinal experiment (MD, n=54; CL, n=54). (d) NMDS of Bray-Curtis dissimilarities between CHC profiles in the single colony experiment (MD, n=59; CL, n=59). Samples are colored by gut microbiota treatment and shapes indicate nurses and foragers in panels (a), (b) and (d). Samples in panel (c) are colored by time of sampling and shapes indicate treatment group.

The independence of CHC profile from microbiota status was confirmed by reanalyzing data from an RNA-sequencing experiment (7) in which we reared CL and MD bees from ten different hives. This experiment included two additional treatments where bees were colonized with either (i) a community of 13 strains covering the predominant species of the honeybee gut microbiota (CL_13; see Supplementary Table 4 in ref. (7)) or (ii) a single core microbiota member, *Bifidobacterium asteroides* (CL_Bifi). Bees from ten different hives were reared in cages of 20 individuals in an incubator for a week after treatment inoculation (one cage per treatment per hive, except for MD bees which were produced in three cages per hive to have a surplus in case of contaminations; see ref. (7) for additional details). To assess the effect of the microbiota on body and gut weight (see next section), we weighed 3-10 bees from 58 cages (548 bees) as well as their guts. We then randomly sampled one to three bees from each of 46 cages (at least one cage per treatment per hive) for CHC analyses (n=120). The bees of the four different treatments differed both in gut microbiota composition and total bacterial load with the MD bees having lower loads than the other three treatment groups, the CL_Bifi bees being dominated by a *Bifidobacterium* phylotype, and the other two colonization treatments having more diverse communities as expected (Extended Figure 4 in ref (7)). CHC analyses of the bees of these four treatments confirmed our previous results (i.e., there was no significant effect of the gut microbiota on the CHC profile; Fig. 2B; PERMANOVA using Bray-Curtis dissimilarities calculated from the centroids of each cage: n=46, *F*(*_3,45_*)=1.13, R^2^=0.07, *P*=0.21).

These results differ from those of Vernier *et al*. (32) who concluded that the honeybee gut microbiota affects the CHC profile of bees. In our experiments, bees were sampled at two time-points in a restricted time-window in the life of adult worker bees (seven and ten days of age for the RNA-sequencing and automated behavioral tracking experiments, respectively). To rule out the possibility that the absence of an effect of the microbiota on the CHC profile could be specific to the two selected time points, we conducted two follow-up experiments. We first reared CL and MD bees originating from nine hives in separate groups of 25 bees (18 cages) and tracked the development of the CHC profiles from day 1 to day 10 post-eclosion by sampling one individual per cage every two days, starting from the day of adult emergence and treatment inoculation (MD, n=54; CL, n=54). We next housed CL and MD bees from a single hive in 20 different cages (ten cages per treatment, which also allowed us to quantify caging effects on the CHC profiles, see below) and sampled them at days 7 (MD, n=30; CL, n=30) and 14 post-emergence (MD, n=29; CL, n=29). While the CHC profiles changed over time, there was again no significant effect of the gut microbiota on CHC profiles in either follow-up experiment (Fig. 2C and D; treatment effects, PERMANOVA using Bray-Curtis dissimilarities calculated from the centroids of each cage: time-series experiment, n=18, *F*(*_1,17_*)=1.22, R^2^=0.06, *P*=0.20; single colony experiment, n=20, *F*(*_1,19_*)=0.85, R^2^=0.05, *P*=0.66; time effects, PERMANOVA using Bray-Curtis dissimilarities: time-series experiment, n=90, *F*(*_1,89_*)=30.30, R^2^=0.21, *P*=0.001; single colony experiment, n=118, *F*(*_1,117_*)=12.09, R^2^=0.08, *P*=0.001).

### The gut microbiota modifies neither body and gut weight nor hypopharyngeal gland size

Because foragers are lighter than nurses (36) and possess degenerated hypopharyngeal (HP) glands (38), we tested whether the microbiota affected these physiological hallmarks of behavioral maturation, using data collected for the RNA-sequencing experiment. There was no significant difference in fresh weight (whole body and gut only) between MD bees and any of the differently colonized bees at seven days of age (Fig. 3A and B; linear mixed effects models fitted by REML with colony of origin and cage as nested random effects: n=548, body weight, *F*(*_3,41_*)=0.94, *P*=0.43, gut weight, *F*(*_3,41_*)=0.22, *P*=0.88). We also measured HP gland size from a subset of the bees (n=28) which were used in brain and gut RNA-sequencing. There was also no significant difference in HP gland size between treatments (Fig. 3C and D; Kruskal-Wallis test: *χ²*=2.75, df=3, *P*=0.43).

**Figure 3.**
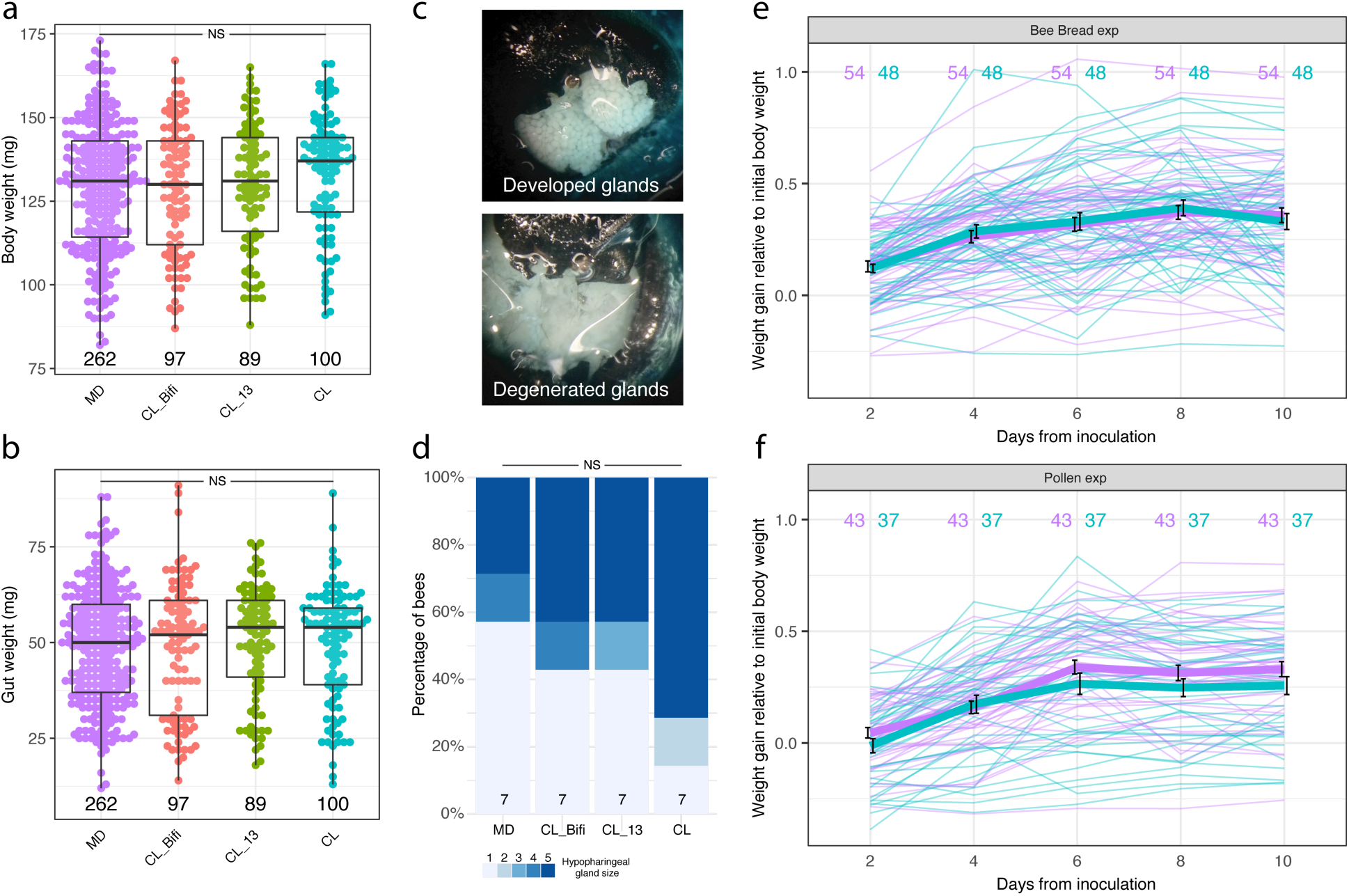
The gut microbiota does not affect weight gain and hypopharyngeal gland size. Boxplots reporting fresh body weight (a) and gut wet weight (b) by gut microbiota treatment group in the RNA-sequencing experiment. Boxplots show the median and first and third quartiles. Whiskers show the extremal values within 1.5x the interquartile ranges above the 75th and below the 25th percentile. (c) Photographs showing examples of maximally developed and degenerated hypopharyngeal glands and (d) proportion of hypopharyngeal gland sizes across gut microbiota treatment groups in the RNA-sequencing experiment. NS, not significant. (e), (f) Fresh body weight growth curves of individual bees colored by gut microbiota treatment group, shown separately for the bee bread (e) and pollen (f) experiments. Values are proportions of initial body weight at the time of adult emergence and gut microbiota colonization. Thicker lines represent mean values and bars indicate SD.

These findings are inconsistent with Zheng *et al.* (31), who found that bees inoculated with a gut homogenate exhibit greater weight gain (for both body and gut weight) than microbiota-depleted bees. However, Zheng *et al.* (31) reported differences in body weight (relative to initial body weight) between CL and MD bees from day 7 onwards, while we had assessed the effect of the gut microbiota on weight only in 7-day-old bees. We also used sterilized pollen to feed the bees, while Zheng *et al.* (31) used a sterilized bee bread diet in their longitudinal experiment. Therefore, we performed two additional experiments to better match the experimental procedure of Zheng *et al.* (31). We reared CL and MD bees from six hives in groups of 30 (one MD and one CL cage per hive). Individuals were uniquely paint-marked and weighed every two days for ten days from the day of treatment inoculation. In a first experiment, bees from three hives were fed bee bread and sugar water *ad libitum*, while in a second experiment bees from three hives were fed pollen instead of bee bread. The microbiota had no significant effect on weight gain in the bee bread experiment and there was no significant interaction between time and treatment either (Fig. 3E; linear mixed effects model fitted by REML with colony of origin, cage and bee individual as nested random effects: n=510, time, *F*(*_4, 400_*)=54.06, *P*<0.0001, treatment, *F*(*_1,2_*)=0.02, *P*=0.91, time*treatment, *F*(*_4, 400_*)=0.66, *P*=0.622). However, there was a statistically significant interaction between time and treatment in the pollen experiment, with MD bees being heavier than CL bees from day 6 onwards (Fig. 3F; linear mixed effects model fitted by REML with bee individual, cage and colony of origin as nested random effects: n=400, time, *F*(*_4, 312_*)=126.98, *P*<0.0001, treatment, *F*(*_1,2_*)=2.10, *P*=0.28, time*treatment, *F*(*_4, 312_*)=2.94, *P*=0.021). This result is in the opposite direction compared to the effect reported by Zheng *et al.* (31) who concluded that the microbiota promotes weight gain.

### Gnotobiotic bees segregate into nurses and foragers with distinct physiology and behavior

While analyzing the CHC profiles of the experiments mentioned above, we realized that bees always clustered into two distinct groups independently of the treatment (Figs. 4A and S1A, B, and C). These two types of CHC profiles corresponded to the typical nurse and forager CHC profiles described in Kather *et al.* (37). To confirm that these CHC clusters represented nurses and foragers, we compared the CHC profiles of our gnotobiotic bees to those of conventional nurses (sampled within hive cells and with pollen-filled guts, n=51) and foragers (sampled on landing boards, carrying pollen and with nectar-filled guts, n=9) from the same ten hives used in the RNA-sequencing experiment. The CHC profiles of these conventional nurses and foragers perfectly segregated into the two clusters (Fig. 4A).

**Figure 4.**
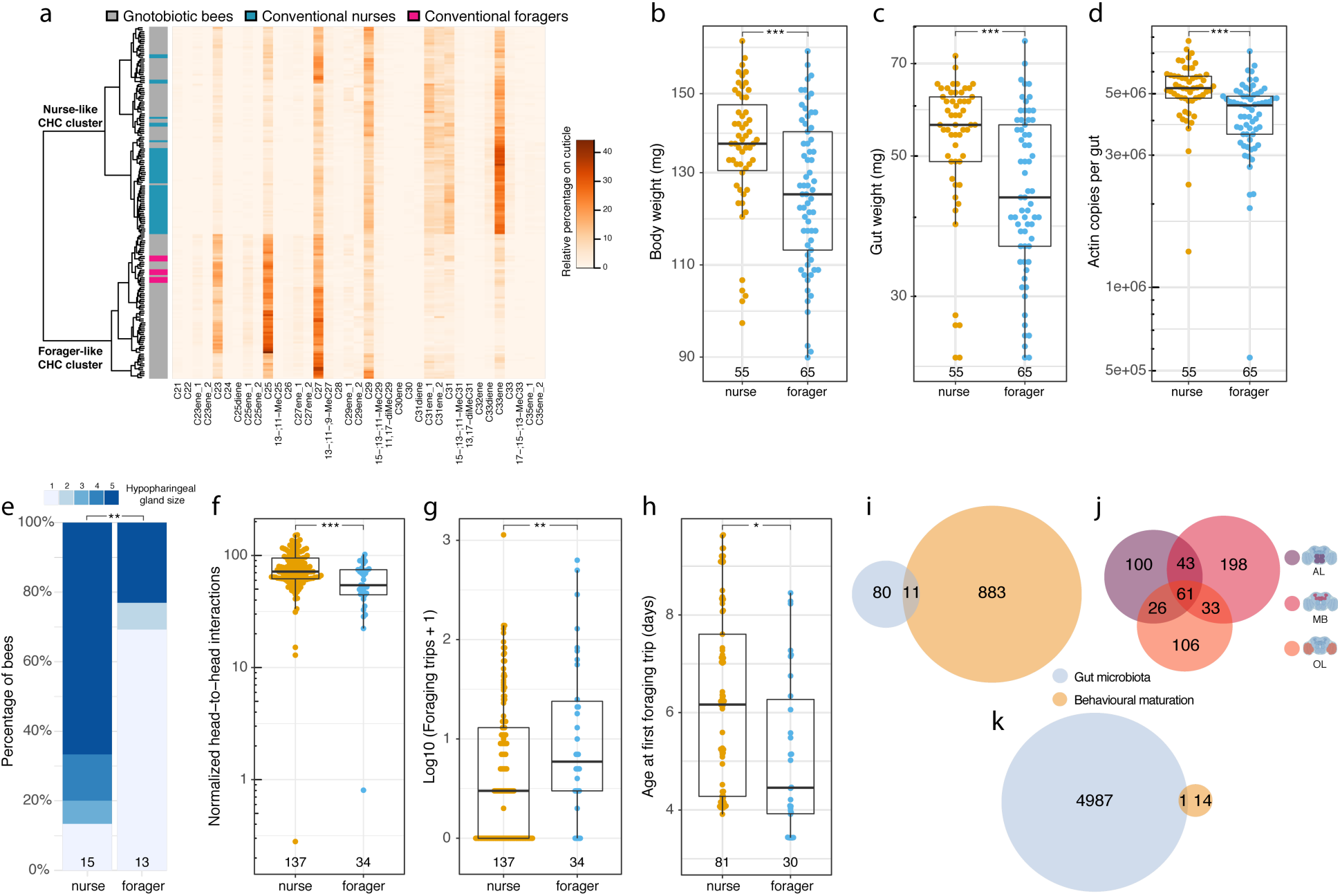
Gnotobiotic bees reared in cages diverge into nurses and foragers showing differences in physiology and behavior. (a) Heatmap of relative abundance of detected CHCs on the cuticle of gnotobiotic bees in the RNA-sequencing experiment (n=120; shown in grey in the annotation column towards the left) and conventional nurses (n=51) and foragers (n=9) collected from the same hives in blue and pink, respectively. The dendrogram towards the left shows clustering of CHC profiles based on Euclidean distances using Ward’s criterion. (b) Fresh body weight (c) gut wet weight, (d) number of *Actin* copies in the gut and (e) hypopharyngeal gland size of CHC-classified nurse and forager gnotobiotic bees in the RNA-sequencing experiment. (f), (g), (h) Boxplots reporting the number of head-to-head interactions (normalized by group size) (f), trips to the foraging arena (g) and the age at first foraging trip (h) of CHC-classified gnotobiotic nurses and foragers in the automated behavioral tracking experiment. \*\*\**P* < 0.001; \*\**P* < 0.01; \**P* < 0.05; NS, not significant. Numbers at the bottom of boxplots and stacked bars in panels (b) to (h) indicate sample sizes. (i) Venn diagram reporting overlap in the brain between the differentially expressed genes (DEGs) associated with the gut microbiota (as identified in (7)) and those associated with behavioral maturation (CHC-classified gnotobiotic nurses *versus* foragers). (j) Venn diagram reporting overlap in DEGs in brain region-specific comparisons of CHC-classified gnotobiotic nurses *versus* foragers. (k) Venn diagram reporting overlap in the gut between the DEGs associated with the gut microbiota (as identified in (7)) and those associated with behavioral maturation. The brain icons were created with BioRender.com.

This CHC-based assignment was further validated by physiological and behavioral data. Consistent with previous studies (36, 38), CHC-classified foragers were lighter than nurses (both for whole body and gut weight) and also exhibited a lower number of *Actin* gene copies in the gut as measured by qPCR on gut DNA extractions, suggesting differences in cell numbers between nurse and forager guts (Fig. 4B, C and D; linear mixed effects models fitted by REML with colony of origin and cage as nested random effects: n=120, whole body weight, *F*(*_1,116_*)=12.61, *P*=0.0006, gut weight, *F*(*_1,118_*)=15.68, *P*=0.0001, log(*Actin* copies), *F*(*_1,110_*)=13.60, *P*=0.0004). Forager-like gnotobiotic bees also had more degenerated HP glands than nurses (Fig. 4E; Kruskal-Wallis test: *χ²*=8.07, df = 1, *P*=0.005). Finally, gnotobiotic bees with a CHC profile typical of foragers interacted significantly less frequently with nestmates, performed more foraging trips, spent more time in the foraging arena and initiated foraging trips earlier than CHC-classified nurses (Fig. 4F, G and H; linear mixed effects models fitted by REML with experimental replicate and sub-colony as nested random effects: social interactions, n=171, *F*(*_1,159_*)=20.17, *P*<0.0001; foraging trips, n=171, *F*(*_1,166_*)=9.18, *P*=0.003, age at first foraging trip, n=111, *F*(*_1,108_*)=6.33, *P*=0.013).

Nurses and foragers are also known to differ substantially in brain gene expression (34). Consistent with this, the comparison of the RNA-sequencing profiles of CHC-classified nurses and foragers revealed a differential expression of 894 genes (i.e., 7% of the transcriptome; Fig. 4I and Supplementary Table 2). To assess whether the gut microbiota affects behavioral maturation-related gene expression, we compared the identity of these genes with those that were differentially expressed as a function of gut microbiota composition (91 genes, ref. (7)). The overlap (11 genes) between these gene lists was not greater than expected by chance (Fig. 4I; hypergeometric test: representation factor = 1.67, *P* = 0.06). Furthermore, differential gene expression by microbiota treatment was most pronounced in the antennal lobe and subaesophageal ganglion region (as shown in (7) for the same experimental bees) while differential gene expression by behavioral maturation was most pronounced in the mushroom body and central complex region (Fig. 4J). Finally, in the gut, 15 genes were differentially expressed between CHC-classified nurses and foragers, of which only one featured among the 4,988 genes differentially expressed between the gut microbiota treatments (Fig. 4K). The overlap between these DEG lists was again not greater than expected by chance (representation factor = 0.16, *P*=0.99). Together these results indicate that, across tissues, the transcriptomic effects of the gut microbiota are not directly related to behavioral maturation.

### Co-housing homogenizes CHC profiles and produces skewed distributions of nurses and foragers

Experiments applying treatments to bees (e.g., microbiota, antibiotics, pesticides) often involve housing bees in shared environments (‘cages’). Co-housing may influence the variables tested due to non-independence (e.g., social interactions) of bees sharing the same cage. We therefore assessed whether uncontrolled co-housing effects on behavioral maturation, which have been previously reported (43), could provide an explanation for inconsistencies between our study and those reporting an effect of the microbiota on host CHC profile and weight gain (31, 32).

We first tested whether co-housing could drive convergence in CHC profiles. To do this, we analyzed the effect of caging on CHC profiles in the experiment where bees from a single hive were placed in 20 different cages (the experimental design involved ten cages per treatment allowing us to assess the effects of caging and microbiota treatment simultaneously). Bees collected from the same cage (5-6 bees per cage) had CHC profiles more similar than bees from different cages (PERMANOVA using Bray-Curtis dissimilarities, n=118, *F*(*_1,117_*)=2.67, R^2^=0.31, *P*=0.001). Additionally, we tested whether the proportions of CHC-classified nurses and foragers were more skewed across cages than expected by chance using the CHC data collected in the automated behavioral tracking experiment because we had CHC data from a minimum of eight bees in each of the 18 cages. This analysis revealed a significant co-housing effect on the proportion of individuals that matured into foragers (range from 0 to 0.6; Chi-square test: *χ²*=30.78, df=17, *P*=0.02).

Because co-housing can lead to skewed proportions of nurses and foragers, individuals within a given cage should not be treated as independent values to study the role of the gut microbiota on behavioral maturation-related phenotypes. Given that previous studies did not control for such an effect, we tested whether the gut microbiota affected the distribution of nurses and foragers across our experiments. For both the 7-and 10-day old bees in the RNA-sequencing and automated behavioral tracking experiments, there was no significant difference in the proportion of nurses and foragers (classified based on CHC profiles) between MD bees and bees of the different colonization treatments (Fig. 5A and B; RNA-sequencing experiment: generalized linear mixed model (GLMM) fitted by maximum likelihood using a binomial distribution with colony of origin and cage as nested random effects, n=120, CL_Bifi, estimate=-0.69, se=0.67, z=-1.02, *P*=0.31, CL_13, estimate=-0.91, se=0.67, z=-1.35, *P*=0.18, CL, estimate=-0.57, se=0.66, z=-0.86, *P*=0.39; automated tracking experiment: GLMM fitted by maximum likelihood using a binomial distribution with experimental replicate and sub-colony as nested random effects, n=177, estimate=0.23, se=0.53, z=0.44, *P*=0.66). This is consistent with the observation that there was no difference between the microbiota treatments in the time bees spent in the foraging arena or the total number of foraging trips performed per bee in the automated tracking experiment. Similarly, there was no significant effect of the gut microbiota on the proportion of foragers at both day 7 and 14 in the experiment designed to assess the effect of co-housing on CHC profiles (Fig 5C; GLMM fitted by maximum likelihood using a binomial distribution with cage as random effect: n=118, time, estimate=-0.87, se=0.58, z=-1.49, *P*=0.14, treatment, estimate=0.72, se=0.72, z=1, *P*=0.32, time*treatment, estimate=-0.74, se=0.84, z=-0.88, *P*=0.38). There was also no significant effect of the gut microbiota on the proportion of foragers at the end (day 10) of either our weight gain experiment with a bee bread diet (Fig. 5D; GLMM fitted by maximum likelihood using a binomial distribution with colony of origin and cage as nested random effects: n=102, estimate=-0.09, se=1.26, z=-0.07, P=0.94), or the weight gain experiment with a pollen diet (Fig. 5E; GLMM fitted by maximum likelihood using a binomial distribution with colony of origin as random effect: n=80, estimate=1.30, se=1.62, z =0.80, *P*=0.42). Finally, the longitudinal CHC experiment, for which we had collected CHC data every two days from adult emergence until day 10 (Fig. 2C), allowed us to more precisely classify the bees that were transitioning between nurse and forager states, as we could identify intermediate groups in the clustering and ordination analyses (Figs. 2C and S1D and E). There was again no statistically significant difference in the proportion of foragers between CL and MD treatments (Fig. 5F; cumulative link mixed model with hive as random effect: n=90, treatment, LR=2.04, *P*=0.15, time*treatment, LR=0.94, *P*=0.33). To further confirm that there was no effect of the gut microbiota on the proportion of foragers, we performed a global analysis comparing the CL and MD treatments across all datasets (n= 602 individuals classified as either nurses or foragers across 94 cages, 35 hives and 6 experiments). For this, we also assessed the effect of the number of co-housed bees at time of sampling. There was a clear effect of time and group size but no effect of gut microbiota treatment on the proportion of foragers, nor an interaction between time and treatment (GLMM fitted by maximum likelihood using a binomial distribution with experiment, colony of origin and cage as nested random effects: n=602, time, estimate=-0.17, se=0.07, z=-2.50, *P*=0.013, group size, estimate=0.02, se=0.01, z=3.35, *P*<0.001, treatment, estimate=0.92, se=0.87, z=1.06, *P*=0.29, time*treatment, estimate=-0.06, se=0.09, z=-0.72, *P*=0.47). These results suggest that the gut microbiota has no effect on the proportion of foragers of honeybee colonies.

**Figure 5.**
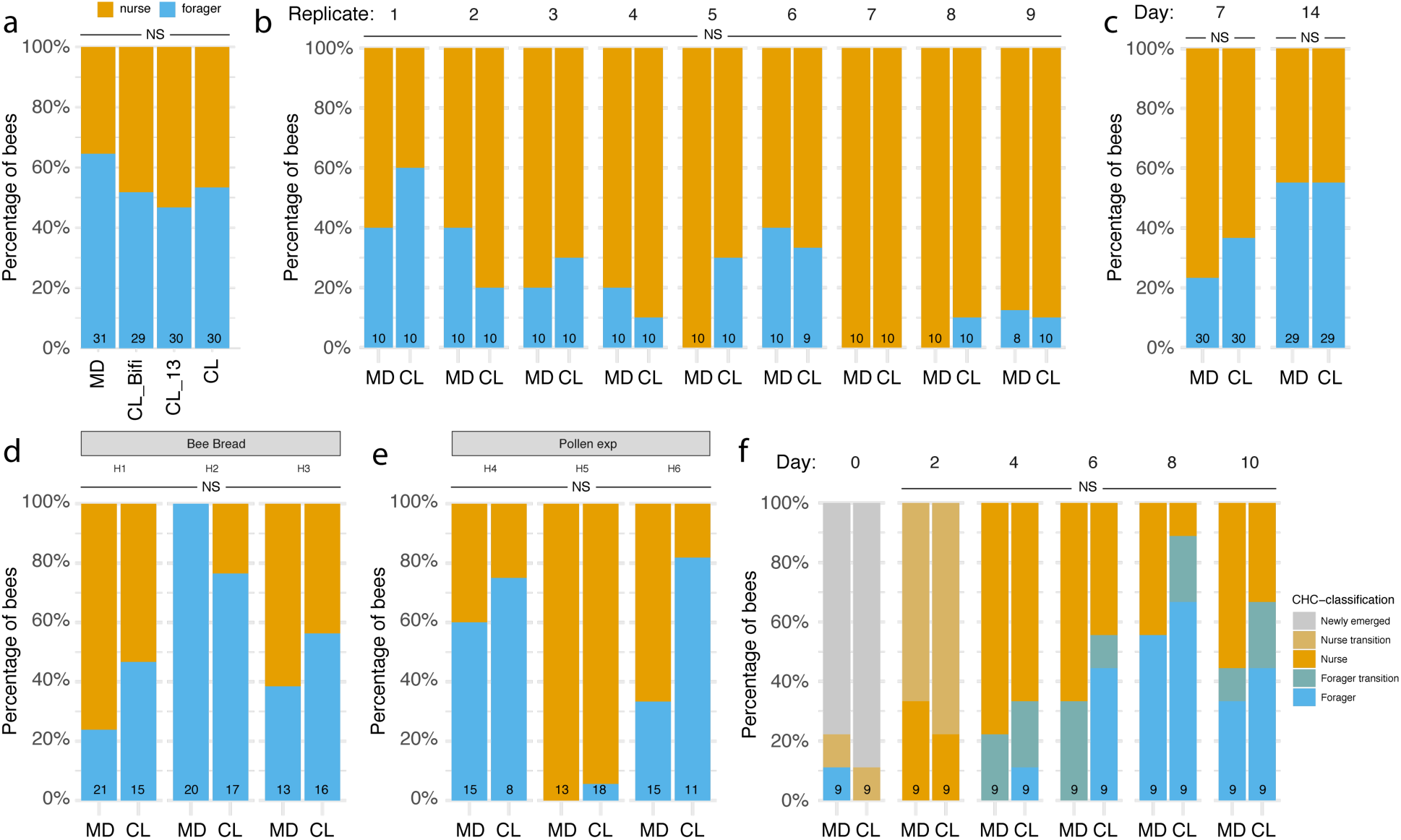
Proportions of nurses and foragers across the experiments. Stacked bars report the percentage of CHC-classified gnotobiotic nurses and foragers (based on clustering of Euclidean distances in CHC profiles using the Ward’s criterion) in the RNA-sequencing (a), automated behavioral tracking (b), single colony CHC (c), longitudinal weight gain with either bee bread (d) or pollen diet (e), and time-series CHC (f) experiments. Numbers at the bottom of stacked bars indicate sample sizes. NS, not significant.

## Discussion

The honeybee is a powerful model to advance evolutionary and mechanistic understanding of host-microbe interactions (26, 44). Previous studies have identified several effects of gut microbes on honeybee phenotypes, including weight (31), CHC profile (32), learning and memory (28, 30) and frequency and patterning of social interactions (7). All these phenotypes change during behavioral maturation (36, 37, 45), with for example foragers being lighter and having different CHC profiles than nurses. This raises the question of whether the reported effects may be indirect (i.e., a consequence of an effect of the microbiota on behavioral maturation). Our experiments showed that while the gut microbiota has a small effect on the time at which bees make their first trip to the foraging arena, there was no effect on the total time bees spent in the foraging arena or the total number of foraging trips performed. Consistent with these behavioral analyses, our data also showed that the microbiota has no significant effect on the proportion of individuals that transition to a forager state and on various physiological hallmarks of behavioral maturation such as CHC profile, gut or body weight, the expression of behavioral-maturation-related genes, or hypopharyngeal gland development. Whether the tendency of colonized honeybees to embark earlier on trips to the foraging arena in the laboratory indicates an effect of the microbiota on the onset of foraging behavior in the field will require further testing.

Our results are in contrast to two previous studies which reported that the honeybee gut microbiota affects CHC profile (32) and promotes weight gain (31). We found that honeybees kept in the same laboratory cage can either take a nurse-like or a forager-like state with correlated changes in physiology and behavior, including CHC profile and body and gut weight. These two types of bees occur in skewed proportions across experimental cages (i.e., individuals within a cage are more similar to each other than individuals between cages). It is likely that previous studies have used only one or few cages per treatment, meaning that the reported associations could stem from co-housing effects. For example, Vernier *et al.* (32) concluded that the honeybee gut microbiota affects the CHC profile of bees. This study involved a series of experiments that identified gut microbiota-associated changes in CHC profile and acceptance behavior of bees. No information is provided on the replication of the cage set-up of the laboratory experiments testing the effects of the gut microbiota on CHC profile. Consequently, it is impossible to determine whether the reported differences in gut microbiota composition and CHC profile in multivariate analyses were due to the experimental treatment or co-housing effects (e.g., social interactions among bees sharing a cage reducing variation in CHC profiles and skewed behavioral maturation producing spurious differences between treatments). However, a re-analysis of the CHC profiles from the key experiment comparing bees inoculated with either live or dead bacterial suspensions showed that, as in our experiments, bees had segregated into nurses and foragers (Fig. S2A) and that there were twice as many foragers in the live inoculum than in the heat-killed inoculum treatment (Heat-killed: 6 foragers and 10 nurses; Live: 11 foragers and 5 nurses; Fig. S2A), driving most of the difference in CHC ordination space (Fig. S2B). Whether the increase in foragers in the live inoculum treatment is due to an effect of the microbiota cannot be determined without details of cage replication. In that respect it should be noted that an effect of the microbiota is unlikely because 16S rRNA gene amplicon sequencing data from this experiment show that bees in both treatment groups had been colonized by core gut microbes and that the microbiota treatments determined a statistically significant difference in the relative proportion of only a few opportunistic bacteria (absolute bacterial loads were not assessed in this study; Fig. S2C and Supplementary Table 3).

Similarly, our results contrast with those of Zheng *et al.* (31) who reported a higher weight gain in microbiota-colonized than microbiota-depleted bees. We could not find such effect across three independent experiments employing larger sample size and cage replication. If anything, in one of our longitudinal experiments there was significant effect of the microbiota in the opposite direction, with CL bees exhibiting reduced weight compared to MD bees from day 6 onwards. However, this effect, unique to one of our three experiments, may have been due to the fact that there were slightly more foragers in the cages assigned to the CL group in this experiment compared to MD cages (Fig. 5E; this difference was not statistically significant). According to Zheng *et al.* (31), bees originated from four hives and were hosted in different cages. Unfortunately, we could not access the original data, precluding testing for co-housing effects. It is still possible that other factors play a role for the discrepancy of these results such as host genotype, gut homogenate used, or seasonal differences between bees.

Our study reveals that bees within cages can be at different stages of their behavioral maturation – a fact that has been previously reported in “single-cohort” colonies (i.e., outdoor hives composed of a few hundred or thousand age-matched young bees (46, 47)), and in groups of age-matched bees kept in the laboratory (43). This had been neglected in studies of the honeybee gut microbiota. Social effects on behavioral maturation can confound gnotobiotic bee experiments and need to be controlled for by randomly sampling individuals from separate cages and increasing the number of replicate cages beyond what has been used in many previous studies. In conclusion, our study indicates that the gut microbiota does not influence the behavioral maturation of honeybees and that previous reports on associations between the gut microbiota and weight gain and CHC profile are likely due to bees within a cage being more similar than between cages because of social interactions.

## Methods

### Rearing of gnotobiotic bees

Across the experiments, bees were reared as previously described (7, 23, 48). Using sterile forceps, we extracted melanized dark-eyed pupae from capped brood cells and placed them in groups of 25-30 into sterilized plastic containers lined with moist cotton. We kept these pupae in an incubator at 70% relative humidity (RH) and 34.5 °C in the dark for 3 days, then transferred newly emerged worker bees into corresponding cup-cages built using a sterile plastic cup placed on top of a 100 mm Petri dish. To colonize bees, an aliquot of a gut homogenate was thawed and diluted 10X in 1X PBS and subsequently 1:1 in sugar water (SW). Microbiota-depleted controls were provided only a 1:1 PBS:SW solution. To inoculate bees, three 100 µl droplets of treatment solution were added to the bottom of each cage. Bees were then kept in their cages in an incubator at 70% RH and 30 °C in the dark (except for the bees in the automated behavioral tracking experiment, which were kept under the tracking systems in groups of ca. 100 bees to monitor their behavior, see below and ref. (7) for additional details), and continuously fed by providing sterile SW and pollen (except for one of the longitudinal weight gain experiments where bee bread was used instead) *ad libitum*.

### Preparation of gut homogenates to inoculate bees

For each experiment, we randomly collected five nurse bees from each of three hives. We anesthetized bees on ice, dissected their guts and placed them individually in 1 mL 1X PBS containing 0.75–1 mm sterile glass beads. Guts were homogenized at 6 ms^−1^ for 45 s using a FastPrep-24 5G homogenizer (MP Biomedicals). The five gut homogenates were pooled by hive of origin and serial dilutions of these pools from 10^−3^ to 10^−12^ were plated onto BHIA, CBA + blood and MRSA + 0.1% L-cys + 2% fructose media using the drop method (10 μl droplets). These plates were then incubated under both anaerobic and microaerobic conditions to verify bacterial growth. Additionally, we prepared lysates of the homogenates by mixing 50 μl of each homogenate with 50 μl lysis buffer, 5 μl proteinase K (20 mg ml^−1^) and 5 μl lysozyme (20 mg ml^−1^) and incubating these mixtures for 10 min at 37 °C, 20 min at 55 °C and 10 min at 95 °C in a PCR machine. Lysates were centrifuged for 5 min at 2,000g and the supernatants used as templates for diagnostic PCR. We performed diagnostic PCRs using specific primers (as done in ref. (7)) to verify the absence of known honeybee pathogens (*Nosema apis*, *Nosema ceranae*, trypanosomatids, *Serratia marcescens*) and fungal growth in bee guts, as well as the presence of bifidobacteria as initial validation that the homogenates contained members of the core gut microbiota. Homogenates with the lowest amplification of pathogen DNA were selected, spiked with glycerol to a final concentration of 20%, and stored at −80°C. Prior to using a selected homogenate in an experiment, we thawed an aliquot and plated it on various media as described above to verify that the homogenates were viable after storage at −80 °C. For the time-series CHC experiment and the two weight gain experiments, we used the same homogenate that had been previously prepared for the RNA-sequencing experiment. The gut homogenates for the automated behavioral tracking experiment and the single colony CHC experiment were prepared anew.

### Measurement of fresh body and gut wet weight

At the end of the RNA-sequencing experiment (see ref. (7) for additional details), we measured fresh body and gut wet weight of the 7-day-old bees across the 58 experimental cages (reared from ten different hives and randomly assigned to four gut microbiota treatment groups). To do this, we anesthetized bees on ice and weighed them using an electric balance sensitive to 0.0001 g. We then dissected their guts as described in ref. (7), placed them in previously weighed 2 mL screw-cap tubes and used the same electric balance to weigh them. The weight of the tube was then subtracted from the total measurement.

Next, we performed two longitudinal weight gain experiments. For each experiment, we reared gnotobiotic bees from three hives in six different cages (one per treatment per hive). Bees from each cage were paint-marked with unique combinations of colors and their body weight was measured every two days for ten days (including the day of adult emergence and treatment inoculation). At each time point, the cages were placed on ice to anesthetize bees and each bee was weighed using an electric balance sensitive to 0.0001 g. At the end of the experiment (day 10) bees were anesthetized on ice, snap-frozen in liquid nitrogen, and stored at −80 °C for subsequent CHC analyses.

### Hypopharyngeal gland size

During brain dissection for RNA-sequencing, we quantified the size of the hypopharyngeal glandular system of 28 bees using a semi-quantitative scale from 1 to 5 (from the most degenerated to the most developed), assigning the score blindly with respect to gut microbiota treatment or CHC group.

### Chemical analysis of cuticular hydrocarbons by GC/MS

We collected cuticular hydrocarbon (CHC) data from bees across multiple experiments. These included the bees at the end of the RNA-sequencing experiment (n=120) and automated behavioral tracking experiment (n=177), when bees were 7 and 10 days old, respectively. We also collected CHC data across the two longitudinal weight gain experiments described above (10-day-old bees; pollen experiment, n=80; bee bread experiment, n=102). We then designed a longitudinal experiment to follow the development of the CHC profile of gnotobiotic bees produced from nine different hives and kept in 18 different cages (one cage per treatment per hive). We collected one bee per cage every two days for CHC analyses starting from the day of adult emergence and treatment inoculation until bees were 10 days of age (n=108). Finally, we performed an additional experiment rearing gnotobiotic bees from a single hive in 20 distinct cages and collecting three bees per cage after 7 and 14 days (n=118). All bees were stored at −80 °C until CHC analyses were performed.

The thorax and abdomen after gut extraction, or only the abdomen for samples of the automated behavioral tracking experiment (thoraxes had been previously used for hemolymph extraction) were submerged in pure hexane for 10 minutes. These extracts were evaporated to a residue of approximately 100 μl. The hexane extracts were run with a DB-5 capillary column (0.25 mm x 30m x 0.25 mm; JW Scientific) on an Agilent 6890-5975 GC-MS at the University of Würzburg (RNA-sequencing experiment), or with an HP-5MS column (0.25 mm x 30m x 0.25 um; Agilent) on an Agilent 8890-5977B GC-MS at the University of Lausanne (all other experiments). A temperature program from 60 °C to 300 °C with 5 °C/min and finally 10 min at 300 °C was used for the RNA-sequencing experiment data, with data collection starting 4 min after injection. The mass spectra were recorded in the electron ionization mode, with an ionization voltage of 70 eV and a source temperature of 230 °C. The chromatography protocol at the University of Lausanne was shortened by ramping the oven from 65 °C to 215 °C at 25 °C/min and then to 300 °C at 8 °C /min. Data were acquired and processed with the ChemStation software v.F.01.03.2357 (Agilent Technologies). Identification of the compounds was accomplished by comparison of library data (NIST 20) with mass spectral data of commercially purchased standards for n-alkanes, diagnostic ions and retention indices.

### CHC data analyses

To calculate the relative abundance of CHC compounds, the area under each compound peak on the GC was quantified through integration using the ChemStation software and divided by the total area under all CHC peaks. The raw data was aligned using the R package GCalignR v.1.0.5 and afterwards analyzed using the packages vegan v.2.6-4 and dendextend v.1.17.1. Polar compounds and contaminations were identified using the mass spectral data (all non-hydrocarbons) and removed from the dataset. Afterwards we removed compounds that were not present in at least half the samples of one treatment or that were only present in trace amounts (<0.1%) in all samples. Lastly, samples which had a too low concentration of CHC compounds (due to failed extractions) were excluded from the analysis. All analyses were done using RStudio v.1.4.1717 and R v.4.1.0 and the package ggplot2 v.3.4.2 for visualization. Area under the peak values were converted to relative proportions, after which we calculated Bray-Curtis dissimilarities between samples and performed non-metric multidimensional scaling (NMDS) ordination analyses and permutational multivariate analysis of variance (PERMANOVA) with 999 permutations to assess differences between experimental groups. To account for sampling multiple individuals from the same cages, we calculated the multivariate centroids from each cage using the *Betadisper* function (package vegan) and tested for the main treatment effect using the resulting matrices, while within-subject effects were tested separately using the original datasets. For PERMANOVA analyses of the CHC profiles in the time-series experiment we removed data from the day of treatment inoculation (day 0) as we did not expect the treatment to have produced immediate effects on CHC profiles (repeating the analysis including day 0 did not change the statistical results qualitatively). Because we analyzed the CHC profiles from the RNA-seq experiment in two separate GC-MS runs, we used the *removeBatchEffect* function in edgeR v.3.34.1 (49) to remove the batch effect prior to plotting the NMDS ordination. We used the *hclust* function of the base R package “stats” to perform hierarchical cluster analyses of Euclidean distances between CHC profiles using the Ward’s criterion prior to plotting heatmaps. Nurses and foragers were then identified based on the resulting clusters. We used generalized linear mixed models fitted by maximum likelihood using a binomial distribution to assess the effect of gut microbiota treatment on the proportion of these CHC-classified nurses and foragers. We always accounted for sampling multiple individuals from the same cages by adding cage as random effect to the models. Based on the hierarchical clustering and ordination analyses of the CHC data collected in the time-series experiment, we were able to identify intermediate clusters (newly emerged bees, bees transitioning to the nurse cluster, nurses, bees transitioning to the forager cluster and foragers). To test the effect of gut microbiota treatment on the proportion of these CHC-clusters, we used a cumulative link mixed model with treatment, time, time*treatment and hive as fixed effects and cage as random effect using the *clmm* function in the package “ordinal” v.2022.11-16. To do this, we again excluded data from day 0 (this did not change the statistical results qualitatively).

### Quantification of foraging tendency under the automated behavioral tracking systems

In the automated behavioral tracking experiment (see ref. (7) for additional details on experimental procedures and data post-processing), bees were housed in a double-box set-up, meaning that they had access to a nest box (kept in constant darkness) connected via a tube to a foraging box (subject to day-night condition cycles). Bees were placed into the nest box at the start of the experiment, allowing us to quantify three metrics for each individual: (i) the time (and hence the age) at which the individual first ventured into the foraging box, (ii) the total proportion of time spent in the foraging arena (i.e., total frames in which an individual was detected in the foraging arena / total number of frames in which the individual was detected in either box), and (iii) the number of box-switches (i.e., each time the individual moved from the nest box to the foraging box and vice versa). We performed all statistical analyses in R v.4.1.0. To assess the effect of the gut microbiota on behavioral variables (average values for each sub-colony) we first checked whether the differences between paired values were normally distributed using the Shapiro-Wilk normality test and then ran either paired *t*-tests or Wilcoxon matched-pairs signed-rank tests.

### RNA-sequencing data analyses

We reanalyzed our previously published RNA-sequencing data (7) to identify differentially expressed genes between CHC-classified nurses and foragers and assess the overlap between these DEGs and those that we had previously identified in gut microbiota treatment comparisons from the same bees (gut, n=38; antennal lobes and suboesophageal ganglion, AL, n=39; mushroom bodies and central complex, MB, n=39; optic lobes, OL, n=38). See ref. (7) for details on data processing to obtain the raw read counts which we reanalyzed in the present study, and for the differential expression analyses of gut and brain between gut microbiota treatment groups. For comparisons of gene expression between CHC-classified nurses and foragers, we used the same parameters as done previously for the gut microbiota comparisons in ref. (7). Briefly, we filtered out genes not represented by at least 20 reads in a single sample using the *filterByExpr* function in edgeR (49). Next, we used the Limma Bioconductor package v.3.48.3 (50) for differential expression analyses. For the gut we used the formula 0 + CHC-classification + batch, whereas for the brain we used the formula 0 + group + batch, where ‘group’ represented every possible combination of brain region and nurse or forager group and ‘batch’ represented the different experimental and RNA-seq library preparation batches. As we had sampled multiple brain regions from the same individuals, we accounted for the individual random effect using the *duplicateCorrelation* function. For the brain, the contrasts between CHC-classified nurses and foragers were performed overall and within each brain region separately. *P* values were adjusted for multiple testing with an FDR of 5%. Hypergeometric tests were used to compare the overlap in DEGs by gut microbiota treatment and by CHC-classification of nurses and foragers in both the gut and the brain.

## Acknowledgments

We would like to thank Christine La Mendola and Catherine Berney for continuous support in the laboratory, Théodora Steiner for assistance with weight measurements and Amelié Cabirol for suggestions on the analyses of foraging rate.

## Funding

University of Lausanne

NCCR microbiomes

The European Union’s Horizon 2020 research and innovation programme under the Marie Skłodowska-Curie grant agreement BRAIN (no. 797113) (JL)

ERC Starting Grant (MicroBeeOme, no. 714804) (PE)

National Centre of Competence in Research, funded by the Swiss National Science Foundation (grant number 180575) (PE)

Swiss National Science Foundation project grant (31003A 160345) (PE)

ERC Advanced Grant (resiliANT, no. 741491) (LKel)

## Author contributions

JL, ETF, TK, PE and LKel conceived and designed the study. JL, PE and LKel acquired funding. PE and LKel supervised the research. JL performed microbiological preparations and gnotobiotic manipulations with assistance from LuK, TK, MMI and AQ. TK performed automated behavioral tracking data analyses with assistance from JL. JL and LuK performed weight measurements. JL performed gut and brain RNA-sequencing analyses. JL and ETF performed CHC extractions. ETF, TS and AQ performed GC-MS runs. ETF, JL and AQ performed CHC data analyses. JL plotted the graphs and performed statistical analyses. JL, TK, LKel and PE drafted the manuscript. All authors contributed to interpreting the data and editing subsequent drafts of the manuscript.

## Data availability

Raw RNA-sequencing data have been deposited in NCBI’s Gene Expression Omnibus and are accessible through GEO Series accession number GSE192784 (https://www.ncbi.nlm.nih.gov/geo/query/acc.cgi?acc=GSE192784), while raw amplicon-sequence data are available on Sequence Read Archive (SRA) under accession PRJNA792398.

## Code availability

Raw data tables, metadata and codes are available on GitHub at https://github.com/JoanitoLiberti/The-honeybee-gut-microbiota-does-not-affect-behavioral-maturation.

## Supplementary Information

**Supplementary Figure 1.** Heatmaps of relative abundance of detected CHCs on the cuticle of gnotobiotic bees in the automated behavioral tracking experiment (a), single colony CHC experiment (b), weight gain experiments (c) and time-series CHC experiment (d). The dendrograms towards the left show clustering of CHC profiles based on Euclidean distances using Ward’s criterion. (e) Non-metric multidimensional scaling (NMDS) of Bray-Curtis dissimilarities between CHC profiles in the time-series CHC experiment, where color represents the CHC clusters identified in the dendrogram in panel (d) and shapes indicate the gut microbiota treatment groups.

**Supplementary Figure 2.** Analyses of the live vs. heat-killed inoculum experiment in Vernier *et al*. (32). (a) Heatmap of relative abundances of detected CHCs. (b) Non-metric multidimensional scaling (NMDS) of Bray-Curtis dissimilarities between CHC profiles, with samples colored either by gut microbiota treatment group or by CHC clusters identified in panel (a). (c) Stacked bars showing the relative abundance of different amplicon sequence variants (ASVs). Sub-bars of the same color show distinct ASVs with the same classification. For ease of visualization, the stacked bars show only ASVs that had a minimum of 2% relative abundance in two samples.

**Supplementary Table 1.** Median relative percentages with median absolute deviation (MAD) of all cuticular hydrocarbons identified in each experiment.

**Supplementary Table 2.** Results of differential gene expression analyses of brain and gut samples between CHC-classified nurses and foragers in the RNA-sequencing experiment, reported in separate sheets for each pair-wise comparison.

**Supplementary Table 3.** ASVs that had an FDR-corrected *P*<0.05 in DESeq2 analyses of differential relative abundance between the live and heat-killed treatments in Vernier *et al*. (32).

