## Supplementary figures and images for "Gut microbiota influences foraging onset without affecting division of labor and associated physiological hallmarks in honeybees"

### Supplementary Figure2

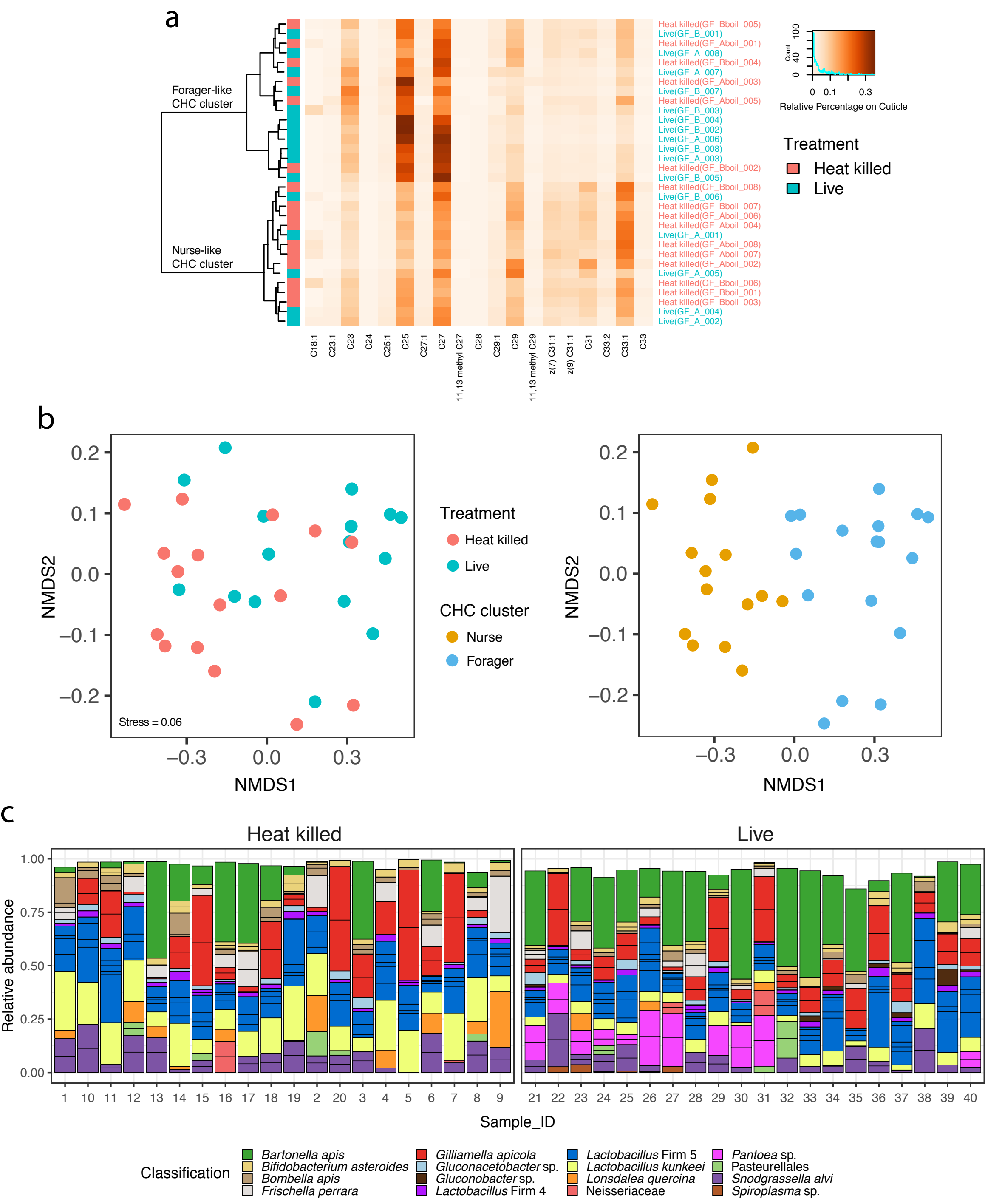

### Supplementary Figure 1

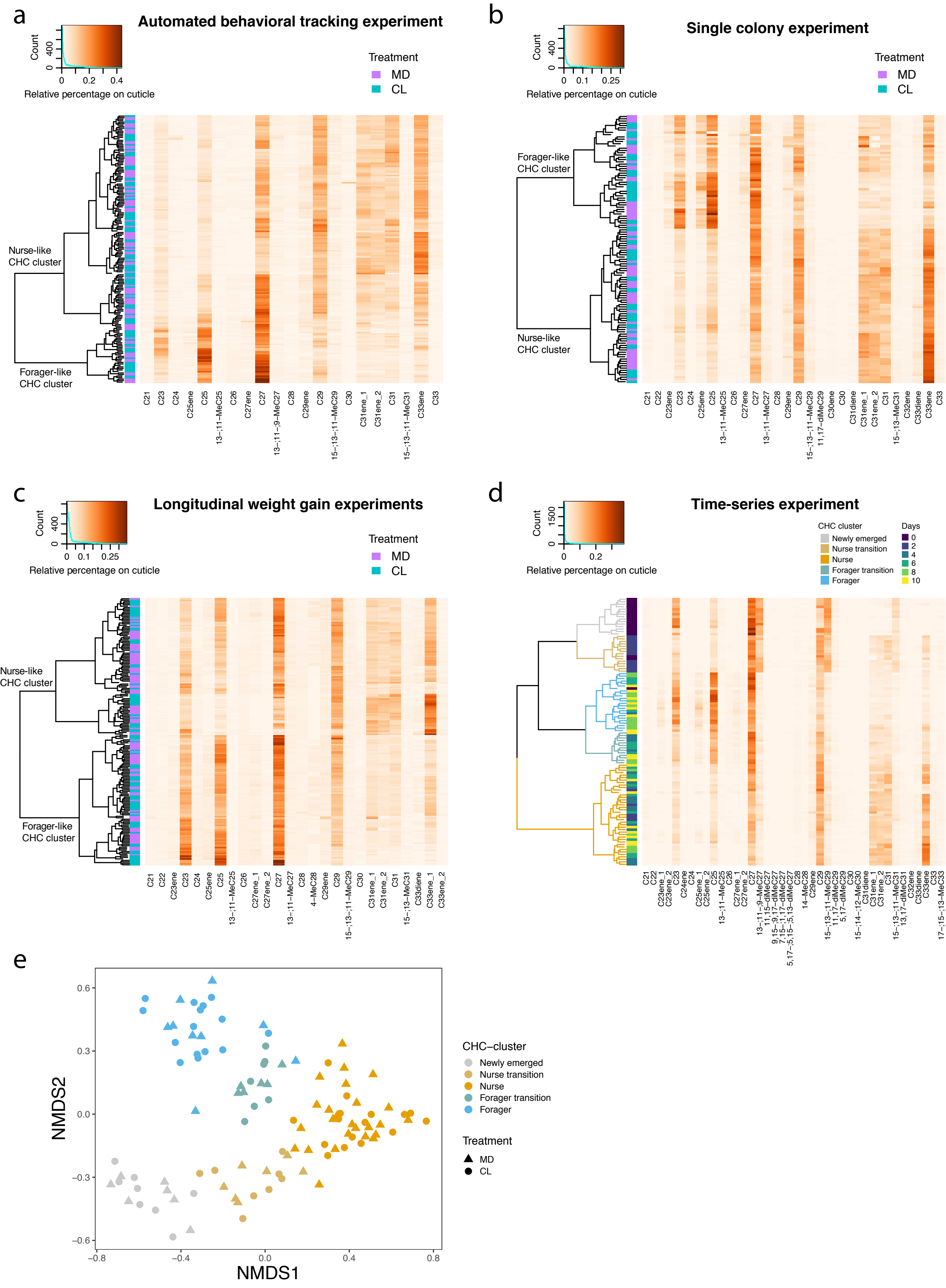
