## Supplementary Table 1 for "Gut microbiota influences foraging onset without affecting division of labor and associated physiological hallmarks in honeybees"

**Supplementary Table 1**. Median relative percentages with median absolute deviation (MAD) of all cuticular hydrocarbons identified in each experiment.

RNA-sequencing experiment:

| Compound | Ret. Index | Hive (n=60) | MD (n=31) | CL_Bifi (n=29) | CL_13 (n=30) | CL (n=30) |
| --- | --- | --- | --- | --- | --- | --- |
| C21 | 2100 | 0.16±0.18 | 0.52±0.24 | 0.52±0.25 | 0.53±0.22 | 0.51±0.17 |
| C22 | 2200 | 0.00 ± 0.08 | 0.11 ± 0.08 | 0.07 ± 0.07 | 0.00 ± 0.08 | 0.09 ± 0.06 |
| C23ene_1 | 2273 | 0.15 ± 0.49 | 0.96 ± 0.89 | 0.59 ± 0.69 | 0.68 ± 0.93 | 0.57 ± 0.71 |
| C23ene_2 | 2279 | 0.00 ± 0.07 | 0.00 ± 0.11 | 0.00 ± 0.06 | 0.00 ± 0.09 | 0.00 ± 0.08 |
| C23 | 2300 | 2.95 ± 3.40 | 10.33 ± 4.61 | 7.68 ± 4.33 | 6.60 ± 4.06 | 6.95 ± 3.97 |
| C24 | 2400 | 0.11 ± 0.18 | 0.47 ± 0.20 | 0.37 ± 0.20 | 0.24 ± 0.22 | 0.31 ± 0.22 |
| C25diene | 2469 | 0.00 ± 0.18 | 0.00 ± 0.14 | 0.07 ± 0.19 | 0.00 ± 0.23 | 0.00 ± 0.14 |
| C25ene_1 | 2473 | 0.42 ± 0.94 | 2.2 ± 1.29 | 1.91 ± 1.16 | 1.27 ± 1.52 | 1.57 ± 1.26 |
| C25ene_2 | 2480 | 0.00 ± 0.16 | 0.26 ± 0.31 | 0.21 ± 0.21 | 0.00 ± 0.20 | 0.17 ± 0.23 |
| C25 | 2500 | 4.61 ± 4.45 | 18.28 ± 7.21 | 11.64 ± 9.47 | 10.15 ± 7.51 | 12.83 ± 8.72 |
| 13-;11-MeC25 | 2534 | 0.16 ± 0.20 | 0.24 ± 0.08 | 0.22 ± 0.10 | 0.28 ± 0.18 | 0.24 ± 0.12 |
| C26 | 2600 | 0.34 ± 0.14 | 0.60 ± 0.20 | 0.69 ± 0.24 | 0.58 ± 0.26 | 0.66 ± 0.31 |
| C27ene_1 | 2673 | 0.18 ± 0.38 | 1.88 ± 0.84 | 1.55 ± 0.77 | 1.02 ± 0.86 | 1.32 ± 0.92 |
| C27ene_2 | 2680 | 0.00 ± 0.13 | 0.42 ± 0.33 | 0.31 ± 0.28 | 0.00 ± 0.28 | 0.33 ± 0.29 |
| C27 | 2700 | 11.44 ± 2.21 | 16.98 ± 3.45 | 20.34 ± 7.03 | 16.19 ± 6.94 | 18.43 ± 6.9 |
| 13-;11-;9-MeC27 | 2731 | 0.98 ± 0.69 | 1.51 ± 0.40 | 1.51 ± 0.35 | 1.81 ± 0.50 | 1.52 ± 0.47 |
| C28 | 2800 | 0.54 ± 0.10 | 0.58 ± 0.14 | 0.57 ± 0.14 | 0.57 ± 0.14 | 0.51 ± 0.11 |
| C29ene_1 | 2875 | 0.00 ± 0.24 | 1.40 ± 1.08 | 1.46 ± 1.06 | 1.69 ± 0.96 | 1.03 ± 084 |
| C29ene_2 | 2879 | 0.91 ± 0.27 | 1.11 ± 1.04 | 0.97 ± 0.97 | 0.59 ± 0.98 | 1.15 ± 0.71 |
| C29 | 2900 | 11.50 ± 1.93 | 9.6 ± 3.97 | 9.82 ± 3.58 | 11.45 ± 4.22 | 10.87 ± 3.64 |
| 15-;13-;11-MeC29 | 2930 | 0.84 ± 0.81 | 1.67 ± 0.36 | 1.58 ± 0.45 | 1.79 ± 0.53 | 1.50 ± 0.45 |
| 11,17-diMeC29 | 2960 | 0.00 ± 0.15 | 0.34 ± 0.18 | 0.00 ± 0.22 | 0.24 ± 0.26 | 0.26 ± 0.21 |
| C30ene | 2981 | 0.00 ± 0.02 | 0.00 ± 0.19 | 0.13 ± 0.17 | 0.00 ± 0.17 | 0.10 ± 0.13 |
| C30 | 3000 | 0.37 ± 0.11 | 0.17 ± 0.13 | 0.24 ± 0.15 | 0.24 ± 0.16 | 0.21 ± 0.12 |
| C31diene | 3060 | 0.42 ± 0.29 | 0.35 ± 0.16 | 0.20 ± 0.18 | 0.21 ± 0.19 | 0.19 ± 0.17 |
| C31ene_1 | 3077 | 7.81 ± 1.71 | 5.72 ± 2.99 | 7.63 ± 4.00 | 7.48 ± 4.13 | 7.19 ± 3.77 |
| C31ene_2 | 3084 | 6.99 ± 1.49 | 3.84 ± 2.02 | 3.95 ± 2.79 | 5.50 ± 2.91 | 5.30 ± 2.91 |
| C31 | 3100 | 10.48 ± 2.25 | 4.00 ± 1.98 | 4.45 ± 2.62 | 5.44 ± 2.35 | 5.30 ± 2.78 |
| 15-;13-;11-MeC31 | 3130 | 0.56 ± 0.48 | 0.97 ± 0.21 | 0.87 ± 0.32 | 1.01 ± 0.39 | 0.84 ± 0.28 |
| 13,17-diMeC31 | 3152 | 0.00 ± 0.15 | 0.22 ± 0.16 | 0.00 ± 0.17 | 0.08 ± 0.18 | 0.16 ± 0.15 |
| C32ene | 3177 | 0.68 ± 0.31 | 0.09 ± 0.26 | 0.00 ± 0.30 | 0.00 ± 0.29 | 0.09 ± 0.27 |
| C33diene | 3257 | 3.52 ± 1.24 | 0.96 ± 0.55 | 0.98 ± 0.71 | 1.02 ± 0.96 | 0.92 ± 0.87 |
| C33ene | 3279 | 25.37 ± 6.02 | 6.98 ± 3.77 | 9.89 ± 6.47 | 8.27 ± 7.21 | 9.70 ± 5.99 |
| C33 | 3300 | 2.33 ± 0.70 | 0.37 ± 0.30 | 0.50 ± 0.44 | 0.39 ± 0.62 | 0.39 ± 0.38 |
| 17-;15-;13-MeC33 | 3327 | 0.00 ± 0.22 | 0.37 ± 0.15 | 0.28 ± 0.22 | 0.29 ± 0.25 | 0.24 ± 0.18 |
| C35ene_1 | 3466 | 0.74 ± 0.31 | 0.00 ± 0.06 | 0.00 ± 0.28 | 0.00 ± 0.34 | 0.00 ± 0.26 |
| C35ene_2 | 3473 | 0.49 ± 0.27 | 0.00 ± 0.01 | 0.00 ± 0.15 | 0.00 ± 0.31 | 0.00 ± 0.11 |

Automated behavioral tracking experiment:

| Compound | Ret. Index | CL (n=90) | MD (n=90) |
| --- | --- | --- | --- |
| C21 | 2100 | 0.53±0.62 | 0.45±0.61 |
| C23 | 2300 | 4.98 ± 2.27 | 4.76 ± 2.23 |
| C24 | 2400 | 0.54 ± 0.44 | 0.49 ± 0.55 |
| C25ene | 2477 | 0.43 ± 0.63 | 0.22 ± 0.33 |
| C25 | 2500 | 6.07 ± 2.66 | 6.41 ± 3.08 |
| 13-;11-MeC25 | 2535 | 0.34 ± 0.51 | 0.30 ± 0.42 |
| C26 | 2600 | 0.72 ± 0.54 | 0.57 ± 0.85 |
| C27ene | 2678 | 0.50 ± 0.74 | 0.27 ± 0.39 |
| C27 | 2700 | 15.97 ± 9.37 | 18.58 ± 9.44 |
| 13-;11-;9-MeC27 | 2733 | 1.32 ± 0.61 | 1.30 ± 0.78 |
| C28 | 2800 | 0.60 ± 0.41 | 0.59 ± 0.44 |
| C29ene | 2882 | 2.55 ± 1.89 | 2.24 ± 1.73 |
| C29 | 2900 | 14.07 ± 2.73 | 14.03 ± 4.28 |
| 15-;13-;11-MeC29 | 2932 | 1.90 ± 1.13 | 1.94 ± 1.02 |
| C30 | 3000 | 0.36 ± 0.54 | 0.00 ± 0.00 |
| C31ene_1 | 3077 | 9.46 ± 3.16 | 8.51 ± 4.41 |
| C31ene_2 | 3084 | 7.60 ± 3.09 | 7.55 ± 3.30 |
| C31 | 3100 | 8.53 ± 4.63 | 9.31 ± 4.39 |
| 15-;13-;11-MeC31 | 3127 | 0.73 ± 0.67 | 0.80 ± 0.77 |
| C33ene | 3276 | 13.34 ± 6.12 | 15.30 ± 5.28 |
| C33 | 3300 | 0.69 ± 1.02 | 0.71 ± 1.05 |

Time-series CHC experiment:

| Compound | Ret. Index | MD (n=54) | CL (n=54) |
| --- | --- | --- | --- |
| C21 | 2100 | 0.49±0.39 | 0.50±0.32 |
| C22 | 2200 | 0.00±0.08 | 0.00±0.09 |
| C23ene_1 | 2273 | 0.17±1.27 | 0.21±1.44 |
| C23ene_2 | 2279 | 0.00±0.19 | 0.00±0.16 |
| C23 | 2300 | 6.34±5.13 | 8.27±5.59 |
| C24ene | 2374 | 0.00±0.07 | 0.00±0.09 |
| C24 | 2400 | 0.06±0.29 | 0.10±0.36 |
| C25ene_1 | 2473 | 0.27±2.05 | 0.35±2.48 |
| C25ene_2 | 2479 | 0.00±0.37 | 0.00±0.51 |
| C25 | 2500 | 6.15±6.51 | 7.25±7.31 |
| 13-;11-MeC25 | 2532 | 0.41±0.32 | 0.34±0.35 |
| C26 | 2600 | 0.39±0.35 | 0.44±0.48 |
| C27ene_1 | 2674 | 0.11±1.04 | 0.28±1.24 |
| C27ene_2 | 2681 | 0.00±0.23 | 0.00±0.43 |
| C27 | 2700 | 17.22±5.87 | 18.67±6.52 |
| 13-;11-;9-MeC27 | 2730 | 3.93±4.71 | 3.33±5.05 |
| 11,15-diMeC27 | 2757 | 0.00±0.20 | 0.00±0.20 |
| 9,15-; 9,17-diMeC27 | 2763 | 0.00±0.12 | 0.00±0.14 |
| 7,15-; 1,17-diMeC27 | 2772 | 0.00±0.08 | 0.00±0.10 |
| 5,17-;5,15-; 5,13-diMeC27 | 2783 | 0.00±0.11 | 0.00±0.11 |
| C28 | 2800 | 0.61±0.29 | 0.56±0.30 |
| 14-MeC28 | 2832 | 0.14±0.26 | 0.09±0.23 |
| C29ene | 2884 | 1.71±1.53 | 1.54±1.12 |
| C29 | 2900 | 12.38±3.97 | 11.74±3.60 |
| 15-;13-;11-MeC29 | 2931 | 4.07±4.89 | 3.42±5.32 |
| 11,17-diMeC29 | 2960 | 0.58±0.74 | 0.51±0.81 |
| 5,17-diMeC29 | 2981 | 0.00±0.27 | 0.00±0.22 |
| C30 | 3000 | 0.00±0.18 | 0.00±0.16 |
| 15-; 14-; 12-MeC30 | 3033 | 0.00±0.09 | 0.00±0.07 |
| C31diene | 3063 | 0.12±0.19 | 0.00±0.15 |
| C31ene_1 | 3076 | 6.01±3.74 | 3.52±3.47 |
| C31ene_2 | 3083 | 5.76±3.19 | 4.20±3.62 |
| C31 | 3100 | 6.26±3.60 | 5.62±4.09 |
| 15-;13-;11-MeC31 | 3129 | 2.27±2.09 | 1.95±2.21 |
| 13,17-diMeC31 | 3157 | 0.41±0.39 | 0.22±0.42 |
| C32ene | 3179 | 0.00±0.28 | 0.00±0.24 |
| C33diene | 3255 | 0.59±0.83 | 0.27±0.70 |
| C33ene | 3276 | 7.06±6.48 | 5.24±6.58 |
| C33 | 3300 | 0.43±0.79 | 0.32±0.80 |
| 17-;15-;13-MeC33 | 3332 | 0.49±0.46 | 0.31±0.51 |

Longitudinal weight gain experiments:

| Compound | Ret. Index | Pollen (n=80) | Bee bread (n=102) |
| --- | --- | --- | --- |
| C21 | 2100 | 0.44±0.41 | 0.61±0.18 |
| C22 | 2200 | 0.56±0.33 | 0.00±0.31 |
| C23ene | 2277 | 0.26±0.73 | 0.99±0.49 |
| C23 | 2300 | 10.07±5.92 | 9.50±3.18 |
| C24 | 2400 | 1.30±0.54 | 0.55±0.56 |
| C25ene | 2476 | 0.65±1.30 | 1.85±0.78 |
| C25 | 2500 | 6.74±7.39 | 13.13±3.56 |
| 13-;11-MeC25 | 2535 | 0.25±0.21 | 0.00±0.15 |
| C26 | 2600 | 1.33±0.44 | 1.08±0.38 |
| C27ene_1 | 2677 | 0.96±1.13 | 1.49±0.71 |
| C27ene_2 | 2684 | 0.00±0.29 | 0.47±0.26 |
| C27 | 2700 | 15.30±7.20 | 18.06±3.39 |
| 13-;11-MeC27 | 2732 | 1.60±0.57 | 1.24±0.52 |
| C28 | 2800 | 1.26±0.28 | 0.77±0.47 |
| 4-MeC28 | 2863 | 0.00±0.17 | 1.90±0.87 |
| C29ene | 2882 | 2.15±1.29 | 2.36±0.56 |
| C29 | 2900 | 12.19±4.28 | 10.94±2.32 |
| 15-;13-;11-MeC29 | 2931 | 1.35±0.58 | 1.11±0.49 |
| C30 | 3000 | 0.94±0.41 | 0.28±0.43 |
| C31ene_1 | 3078 | 6.68±3.73 | 5.51±1.58 |
| C31ene_2 | 3084 | 5.18±3.55 | 4.27±1.54 |
| C31 | 3100 | 5.92±3.66 | 5.36±1.62 |
| 15-;13-MeC31 | 3133 | 0.62±0.43 | 0.51±0.29 |
| C33diene | 3252 | 0.72±1.11 | 1.17±0.84 |
| C33ene_1 | 3268 | 7.20±8.24 | 9.33±3.00 |
| C33ene_2 | 3284 | 0.59±0.78 | 0.38±0.30 |

Single colony experiment:

| Compound | Ret. Index | CL (59) | MD (59) |
| --- | --- | --- | --- |
| C21 | 2100 | 0.52±0.29 | 0.45±0.27 |
| C22 | 2200 | 0.14±0.12 | 0.11±0.07 |
| C23ene | 2276 | 1.22±1.79 | 0.58±1.37 |
| C23 | 2300 | 6.13±4.27 | 5.44±4.64 |
| C24 | 2400 | 0.28±0.37 | 0.18±0.33 |
| C25ene | 2476 | 3.81±3.65 | 1.65±3.17 |
| C25 | 2500 | 8.85±6.53 | 6.91±6.82 |
| 13-;11-MeC25 | 2534 | 0.28±0.11 | 0.26±0.11 |
| C26 | 2600 | 0.52±0.47 | 0.41±0.29 |
| C27ene | 2678 | 1.70±1.73 | 1.02±1.27 |
| C27 | 2700 | 14.42±4.40 | 13.16±4.30 |
| 13-;11-MeC27 | 2733 | 1.60±0.63 | 1.52±0.79 |
| C28 | 2800 | 0.51±0.18 | 0.50±0.16 |
| C29ene | 2883 | 2.31±1.31 | 1.99±1.58 |
| C29 | 2900 | 11.44±3.24 | 12.25±3.21 |
| 15-;13-;11-MeC29 | 2932 | 1.48±0.65 | 1.45±0.77 |
| 11,17-diMeC29 | 2961 | 0.19±0.18 | 0.13±0.20 |
| C30ene | 2981 | 0.24±0.18 | 0.29±0.19 |
| C30 | 3000 | 0.18±0.11 | 0.21±0.14 |
| C31diene | 3063 | 0.26±0.22 | 0.23±0.21 |
| C31ene_1 | 3076 | 9.04±3.32 | 9.19±3.89 |
| C31ene_2 | 3084 | 7.68±3.09 | 8.42±3.00 |
| C31 | 3100 | 6.65±2.52 | 7.78±3.21 |
| 15-;13-MeC31 | 3132 | 0.64±0.35 | 0.68±0.43 |
| C32ene | 3160 | 0.28±0.24 | 0.33±0.22 |
| C33diene | 3248 | 1.22±1.20 | 1.48±1.03 |
| C33ene | 3263 | 14.17±5.99 | 15.80±5.62 |
| C33 | 3300 | 0.44±0.43 | 0.72±0.64 |
